# Positional cues, not Notch, direct Neuroblast selection during early neurogenesis in the *Drosophila* embryo

**DOI:** 10.64898/2026.03.30.715196

**Authors:** David Green, Khalil Mazouni, Marcel Nos, Francois Schweisguth

## Abstract

Notch-mediated lateral inhibition is a conserved patterning process that controls alternative cell fate decisions and produces regular cell fate patterns. Prevailing models posit that lateral inhibition singles-out cells from fields of initially equipotent cells by amplifying stochastic fluctuations of Notch or pre-existing fate biases. Here, we revisited the role of Notch in early *Drosophila* neurogenesis, studying the dynamics of Neuroblast specification by live imaging the transcription of two proneural genes, *scute* and *lethal of scute*. We found that proneural gene expression is biased spatially along the dorsal-ventral axis prior to germ band extension and that early proneural expression predicts Neuroblast fate acquisition. This indicated that Neuroblast specification is pre-patterned by positional cues. Additionally, positional cues appeared to instruct individual cells to delaminate in a correct stereotyped pattern in proneural mutant embryos. Finally, contrary to current models, Notch signaling, measured by *E(spl)m8* expression, was not detectable within proneural clusters until after Neuroblasts had initiated delamination. This indicated that Notch functions to stabilize rather than initiate fate decisions. We therefore propose that positional cues, not Notch, single-out Neuroblasts during early *Drosophila* neurogenesis, challenging long-held assumptions about the role of Notch in Neuroblast selection.

## Introduction

A fundamental question in developmental biology is how cells acquire distinct fates to generate complex tissues with stereotyped cell fate patterns. While patterns can arise from a cascade of positional cues that establish a series of subdivisions^1^ or via self-organized processes^2,3^, it has become increasingly clear that many patterns are produced by a combination of self-organization and positional information-based processes^4,5^. Lateral inhibition is a conserved patterning process whereby equipotent cells inhibit each other from adopting the same fate via cell-cell interactions mediated by the membrane-bound ligand Delta and its receptor Notch^6,7^. In the absence of positional cues lateral inhibition produces a salt and pepper pattern of cell fates^8,9^. Genetic studies and mathematical models have indicated that binary fate decisions rely on the amplification of stochastic fluctuations in Notch via an intercellular feedback loop^10–12^. Self-organization by Notch can also be guided by positional cues to produce stereotyped patterns of cell fates^4,13^. During early neurogenesis of the *Drosophila* embryo, Notch is required to produce a stereotyped pattern of Neuroblasts (NBs). How positional information integrates with Notch signaling to establish this stereotyped pattern is still poorly understood.

During early neurogenesis, the ventral neuroectoderm (VN) of the *Drosophila* embryo produces a fixed number of NBs that delaminate in five successive waves, noted S1-S5^14,15^. These NBs form an orthogonal array organized along three columns along the dorsal-ventral axis (DV) and four rows along the anterior-posterior axis (AP) (Fig 1A,B)^15,16^. Each of these NBs emerge from small groups of 5-6 cells expressing proneural genes of the *Achaete-Scute* complex (*AS-C*)^17–21^. In the first wave of neurogenesis (S1) these ProNeural Clusters (PNCs) are positioned along the AP and DV axes by upstream patterning genes, generating an orthogonal array of ∼10 PNCs which produces 10 unique NBs per hemisegment^16,17,22,23^. Within each PNC, inhibitory cell-cell interactions mediated by Delta-Notch signaling is required to select a single cell to become a NB^17,20,21,24,25^, which then constricts its apical surface and delaminates from the epithelium^26,27^. Interestingly, NBs emerge during germband extension (GBE), when cells of the VN undergo neighbor exchanges to drive the convergence-extension of the VN^28^. Thus, PNCs appearing as compact groups of 5-6 cells at the end of GBE are predicted to arise from groups of cells that are more elongated along the DV axis prior to GBE (Fig 1C). Considering this, the current view proposing that positional cues act within PNCs to produce a fate bias that is then amplified by Notch to select NBs raises many questions: How are positional cues maintained throughout GBE? Alternatively, are positional cues interpreted prior to GBE? If so, are NBs selected prior to GBE? And when does Notch select NBs relative to neighbor exchanges?

**Figure 1.**
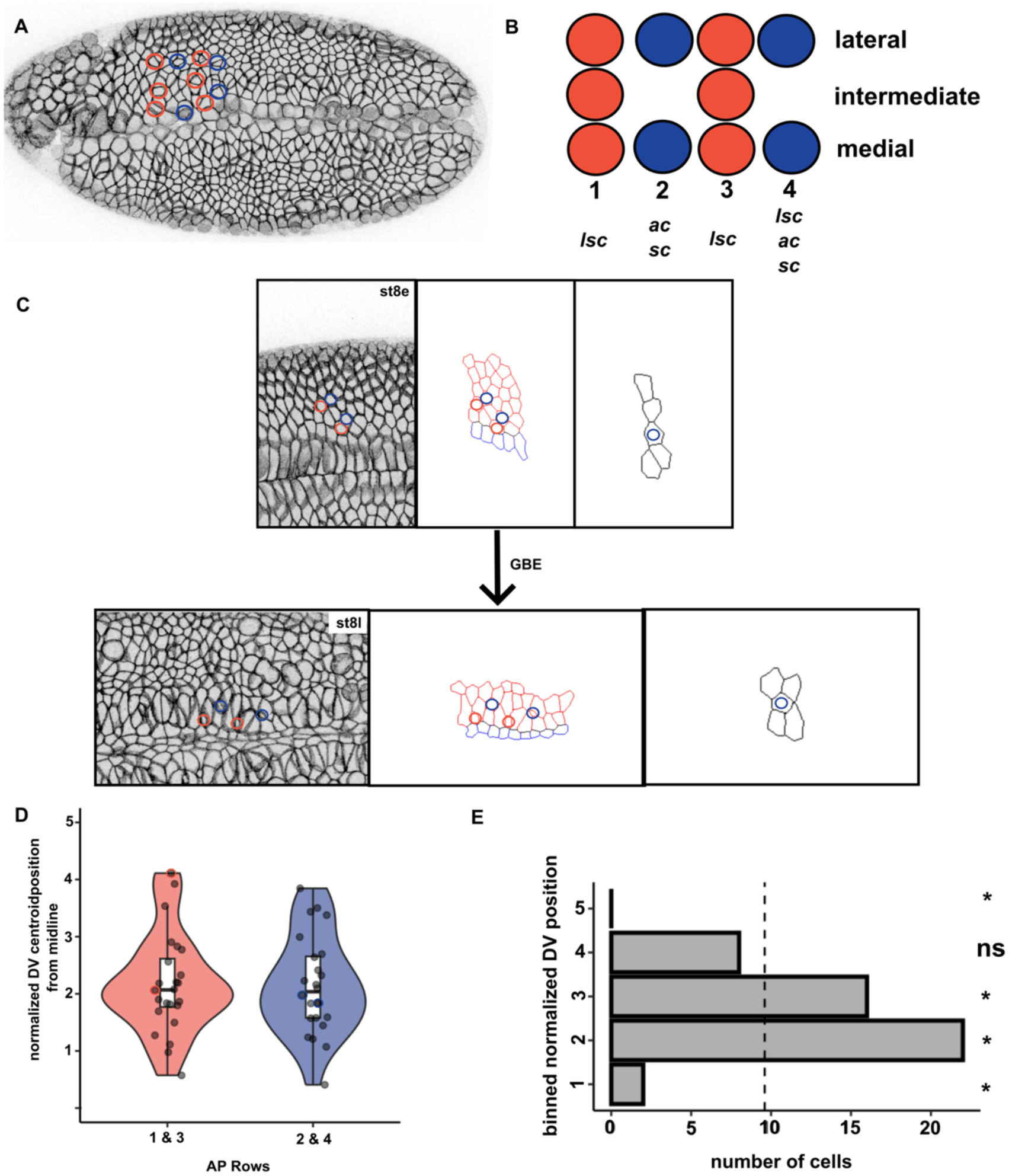
Medial NBs occupy specific DV positions prior to GBE (A,B) Ventral view of a PHmCh embryo at late stage 8 (A), outlining the segmental pattern of S1 NBs (schematized in B). The S1 NBs are organized into four AP columns, each defined by a combination of proneural factors^16^ (NBs from columns 1 and 3 in red; from columns 2 and 4 in blue) and three DV rows, defined by the expression of the Vnd (medial row), Ind (intermediate row) and Msh factors (lateral row)^16^. (C) Snapshots from a movie of a PHmCh embryo showing the VN before (top) and shortly after (bottom) the onset of GBE. VN cells from one segment are outlined in red (mesectoderm, blue). A single AP column, containing a delaminating NB (top-right), rearrange to form a more compact group of cells (bottom-right). Small circles marked NBs from AP (color code like in B). (D) Plots showing the DV distribution of the pre-GBE position of presumptive NBs for two sets of columns (1,3 and 2,4). Violin plots show density distribution; box plots show median and quartiles. Individual points represent single NBs. (E) Chi-square analysis reveals non-uniform DV distribution (X^2^=19.3, degrees of freedom, df=3, p-value, p<0.001); stars indicate positions with standard deviation (sd) residual >3.33 or <-3.33. Dashed line indicates expected frequency under uniform distribution.

Here, we investigated the dynamics of NB fate specification in the medial VN using live imaging. We first found that cells becoming medial S1 NBs occupied a biased position along the DV axis prior to GBE. Using functional MS2 knock-in alleles of the proneural genes *scute* (*sc*) and *lethal of scute* (*lsc*), we identified a DV pattern of proneural expression prior to GBE which predicted NB fate, indicating that NB specification is indeed pre-patterned by positional cues. Proneural genes were required for the timely onset of apical constriction but were dispensable for NB delamination, suggesting that positional cues may act in parallel to proneural genes to select individual delaminating cells in a correct stereotyped pattern. Finally, we found that the Notch effector Enhancer-of-split m8 factor [E(spl)m8] was not detected until after NBs begin to apically constrict, indicating that Notch does not act to single-out NBs in the early embryo. These findings challenge the current model of NB specification and led us to propose that Notch acts after the onset of NB delamination to stabilize and reinforce an initial fate difference that results from an early proneural bias established prior to GBE.

## Results

### Future medial NBs occupy biased DV position prior to GBE

NB delamination occurs during GBE, a period of dramatic tissue remodeling during which VN cells undergo neighbor exchanges through T1 transitions^28–30^. Expectedly, these cell-cell intercalations change the geometry of the cell clusters corresponding to the presumptive PNCs. The live imaging of embryos expressing the PH domain of PLCγ fused to Cherry (PHmCh) as a fluorescent membrane marker showed that rows of cells oriented along the DV axis were reorganized into more compact clusters corresponding to presumptive PNCs as embryos elongated along the AP axis (Fig 1C). While delaminating NBs are found at stereotyped positions, the position of the future NBs relative to other PNC cells prior to GBE has not been examined. Here, we first addressed whether NBs occupy a biased position relative to their PNC neighbors prior to GBE.

To precisely map the spatial origin of NBs, we imaged living embryos expressing PHmCh. NBs were identified in stage 9 embryos based on their delamination and were then backtracked to determine their position at stage 7, prior to GBE (Fig 1C). We focused on medial NBs since these cells remained in the field of imaging over stages 7-9, i.e. throughout GBE. Using NBs from the intermediate column as landmarks (Fig 1C), we were able to identify NBs from rows 1/3 and 2/4 (Fig 1A-C, Movie 1). After back-tracking individual medial NBs across multiple segments, we measured their DV position relative to the mesectoderm prior to GBE. This indicated that most NBs appeared to originate from a position located two cells away from the mesectoderm (Fig 1D). To test whether NBs arise from non-random positions, we binned normalized DV position into five cell lengths and performed statistical analysis on these distributions. This showed that positions 1 and 5 were significantly under-represented whereas position 2 and 3 were significantly over-represented (Fig 1E). Thus, medial NBs do not arise from random pre-GBE position but instead occupy spatially biased positions along the DV axis. We therefore conclude that NB selection appears to be biased by early DV cues.

### Live imaging of proneural gene transcription revealed an early DV bias

Since proneural factors are expressed in the VN prior to GBE^18,19^, we wondered whether the observed DV bias in the position of the future NBs might result from an early DV pattern of proneural gene expression. Recent advances in MS2/MCP-based live imaging of transcription have enabled real-time tracking of gene expression with high spatial-temporal resolution^31–34^. The interaction of the MS2 bacteriophage coat protein (MCP), fused to a fluorescent protein, with multiple repeats of a stem-loop structure from the phage genome (MS2 repeats) present within nascent transcripts creates fluorescent foci at sites of active transcription in living cells and embryos^31,32^. To visualize proneural gene expression dynamics, we therefore inserted 24xMS2 stem loop repeats into the 5’ UTR of *sc* and *lsc* using CRISPR-mediated homologous recombination, generating *ms2:sc* and *ms2:lsc* lines (Fig 2A). Both lines were homozygous viable with no observable developmental defects, suggesting that MS2 insertion did not disrupt normal proneural function. The live imaging of *ms2:sc* and *ms2:lsc* embryos with maternally deposited MCP-GFP revealed bright fluorescent puncta throughout the VN (Fig 2B, Movies 2,3). By segmenting the MS2 foci (using MCP-GFP) and tracking both cells (apically using PHmCh) and nuclei (using His2A-RFP), we were able to quantify proneural transcriptional activity in individual cells over time (see Methods for segmentation and tracking details). These datasets were first used to ask when the *sc* and *lsc* genes were transcribed in the VN relative to GBE. Transcriptional foci were first detected 10 minutes (min) before GBE in *ms2:lsc* (Fig 2C) and 5 min before GBE in *ms2:sc* embryos (Fig 2C’; onset of GBE is t=0 in Fig 2), consistent with published expression data^20,21^. To next determine whether proneural activity showed spatial bias along the DV axis, we tracked medial VN cells and measured the MCP-GFP signal at transcription foci. We observed that neighboring cells within the same row of cells exhibited differences in *lsc* and *sc* gene expression by the onset of GBE (Fig 2D, Movies 4,5). To determine whether this early expression heterogeneity showed a spatial pattern, we mapped the distribution of *lsc* and *sc* expression along the DV axis 10 min prior to the GBE onset (Fig 2E,F). Pseudo-coloring cells by their cumulative signal intensity revealed clear spatial patterning within the medial VN. High-expressing cells were not uniformly distributed but instead showed enrichment at intermediate DV positions. Quantification of proneural expression revealed a biased distribution along the DV axis for both the *sc* and *lsc* genes, with peak expression occurring 2-3 cell diameters from the mesectoderm (Fig 2E’,F’). This expression bias matched the spatial bias in the position of the future NBs (Fig 2E’’,F’’). Of note, the accumulation pattern of the Lsc protein, studied here using an endogenous GFP-tagged version of Lsc (see Methods), showed a similar correlation (Fig S1; note that GFP-Lsc was not detectable by live imaging, presumably due to its fast turnover relative to the maturation kinetics of eGFP). Together, our data showed that the early pattern of expression of the *lsc* and *sc* genes correlated with the NB fate bias observed along the DV axis prior to GBE (Fig 1E).

**Figure 2.**
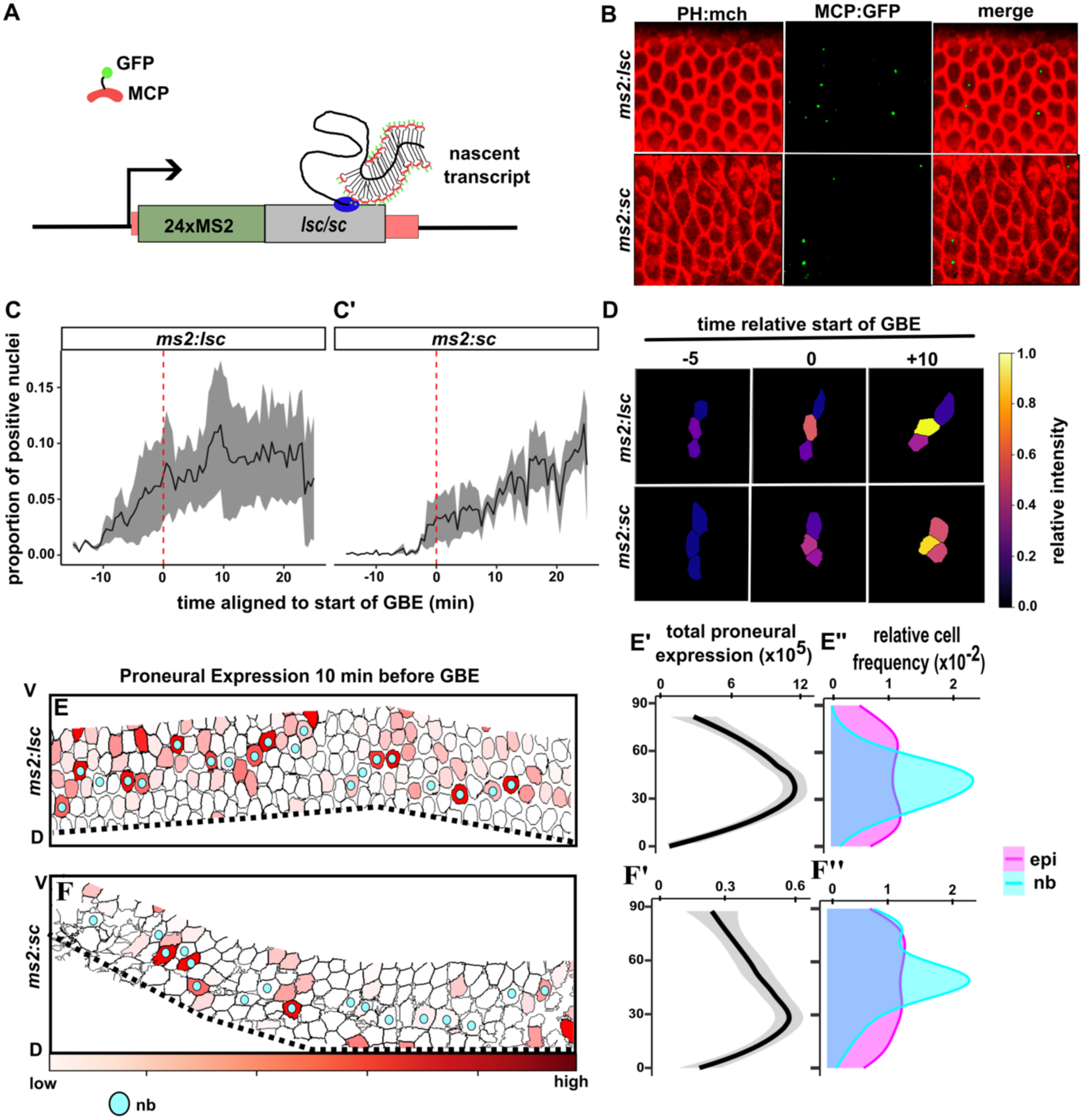
Live imaging revealed early proneural transcription bias (A) Schematic of the MS2/MCP system. Nascent *lsc* (or *sc*) transcripts carrying 24xMS2 stem loop repeats in their 5’ UTR interact with MCP-GFP to produce fluorescent foci. (B) Representative images of a stage 8 embryo expressing the *ms2:lsc* (top) and *ms2:sc* (bottom) genes (PHmCh, red; MCP-GFP, green). (C,C’) Plots showing the fraction of cells with transcriptional foci over time, aligned to GBE onset (t=0, red dashed line; n≈ 600 VN cells from 3 embryos per genotype). Line, mean; shaded area, sd. (D) Time-lapse montage of transcription dynamics in cells from the same DV column which were pseudo-colored by cumulative MS2 spot intensity. Time relative to GBE in min. (E-F’’) Spatial distribution of *lsc* and *sc* transcription before GBE in representative embryos (NBs are shown with blue dots), with VN cells pseudo-colored by total cumulative MS2 intensity over a [-10,0] period (in min) before GBE (E,F; black dashed line, mesectoderm boundary). The cumulative expression of the *lsc* and *sc* genes along the DV axis (E’,F’) and the relative frequency of cells fated to become NBs (E’’,F’’; cyan; all other VN cells in magenta) showed that proneural transcription prior to GBE correlated with NB fate acquisition during GBE. X-axis represents the position along the DV axis, with the mesectodermal boundary set as 0. Grey shaded region represents 95% confidence interval. n=4 embryos, ∼ 30 NBs and ∼100 other VN cells (noted epi here) per genotype.

### Early proneural expression predicts NB fate

These correlations suggested that early proneural activity prior to GBE may act to bias NB fate selection during GBE. To test this, we measured the level of *sc* and *lsc* expression prior to GBE in each medial VN cells and asked whether these values correlated with fate acquisition during GBE. This analysis indicated that presumptive NBs, identified by their later delamination, had significantly higher levels of *lsc* and *sc* expression than their neighbors (Fig 3A,A’). This supported the view that heterogeneities in early proneural expression could bias NB fate selection during GBE. To further test whether early proneural gene expression correctly predicts cell fate, we performed a logistic regression analysis that relates proneural activity 10 min before GBE to NB fate adoption (Fig 3B,B’). For both *lsc* and *sc*, the logistic-regression curves showed that cells with high early proneural expression had increased probability of becoming NBs (7- and 14-fold increase of odds for each standard deviation increase in *lsc* and *sc* intensities, respectively). Thus, early proneural expression strongly predicted NB fate acquisition. This predictive power is striking given that intensity values were measured prior to GBE, i.e. shortly after the onset of detectable proneural expression. We therefore conclude that an early fate bias regulates the NB fate decision prior to the GBE onset.

**Figure 3.**
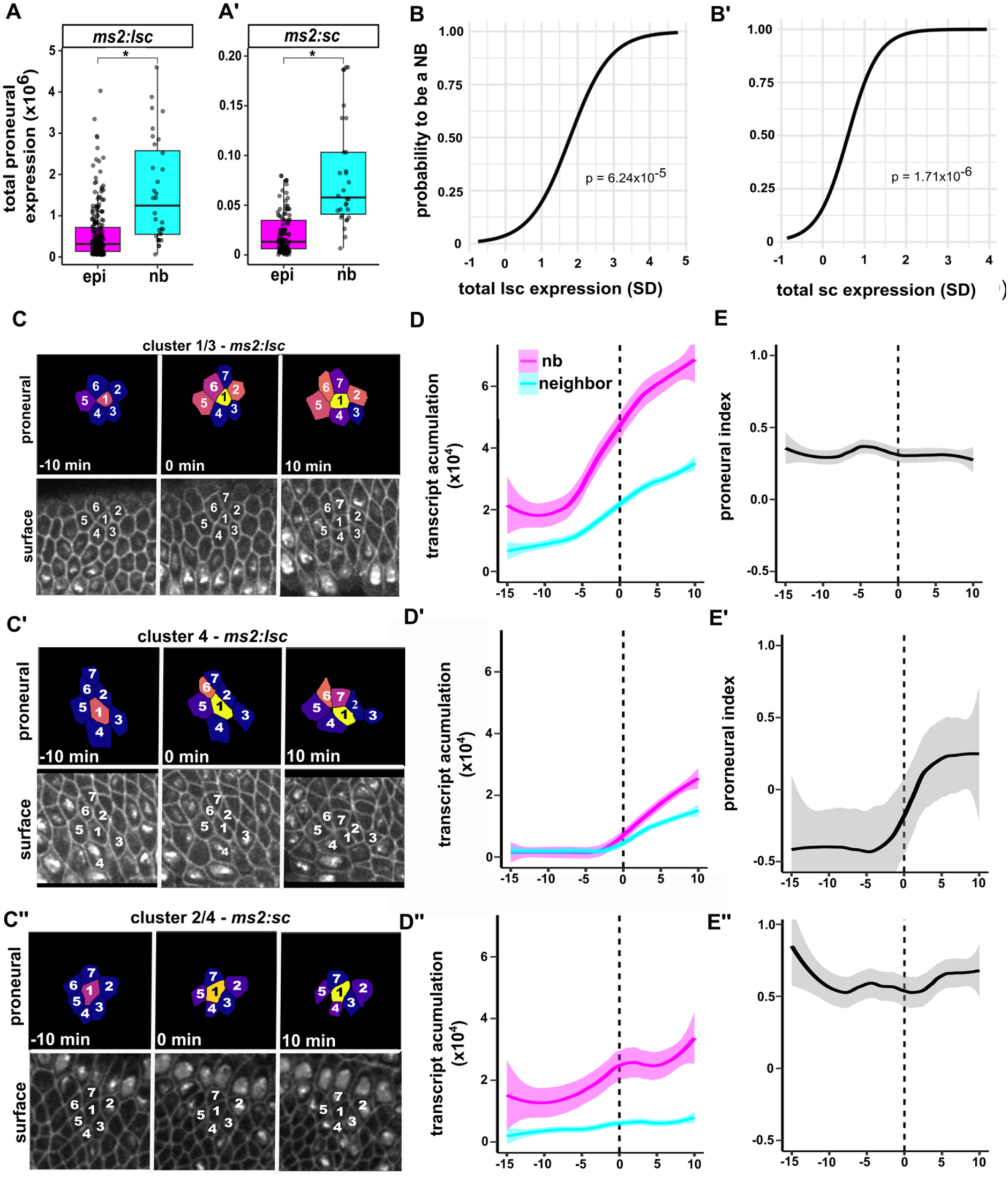
Early proneural expression predicts NB fate acquisition (A,A’) Presumptive NBs showed higher levels of *lsc* and *sc* transcription prior to GBE than other VN cells, noted her as epi (Wilcox test, p< 0.05). Box plots show median and quartiles. (B,B’) Logistic regression analysis relating early proneural gene expression to eventual NB fate. Curves show probability of adopting the NB fate as a function of proneural expression levels. Early proneural expression was a significant predictor of cell fate (chi-squared test based on regression analysis). (C-C’’) Time-lapse montage showing proneural expression dynamics (*ms2:lsc* in C,C’; *ms2:sc* in C’’) within individual PNCs corresponding to different AP columns, expressing different combination of proneural genes (see Fig 1B). In top rows, cells are numbered around future NB (cell 1) and pseudo-colored by cumulative MS2 intensity (as in Fig 2D; t=0, onset of NB delamination). Bottom rows show apical surface (PH:mCh). (D-D’’) Average transcription dynamics in different PNCs, showing the cumulative expression over time of *lsc* (D,D’) and *sc* (D’’) aligned to the start of NB delamination. (E-E’’) Plots showing the evolution of proneural index values (measuring the relative transcription of proneural genes in NBs compared to their neighbors) over time. Grey region represents 95% confidence interval. n=8 NBs/PNCs per PNC category.

Different PNCs express different proneural genes (Fig 1B; all S1 NBs can be studied using the *lsc* and *sc* genes). To better understand the expression dynamics within individual PNCs, we studied the expression dynamics of *sc* and *lsc* separately in clusters 1 and 3 (*lsc* only), 2 (*ac* and *sc*) and 4 (*lsc*, *ac* and *sc*) by tracking individual NBs and their immediate neighboring cells (defined by apical surface contact; Fig 3C-C’’). In clusters 1 and 3, the *lsc* gene was expressed early, before the onset of apical constriction (t=0 in Fig 3), and presumptive NBs exhibited significantly higher expression than the neighboring cells from the earliest time points (Fig 3D). While *lsc* gene expression remained high in NBs, expression increased only gradually in the neighboring cells, leading to a persistent difference in expression levels (Fig 3D). To quantify differences in proneural activity within PNCs, we defined a “proneural index” that measures the relative expression difference in NB and neighboring cells (see Methods). The values for this index remained relatively constant over time in clusters 1 and 3 (Fig 3E). This indicated that the initial bias in proneural expression was maintained until NBs delaminate. In contrast, the expression dynamics of the *lsc* (in clusters 4) and *sc* genes (in clusters 2 and 4) revealed differences in early proneural expression. In clusters 2 and 4, early *sc* expression levels were higher in presumptive NBs than in their neighboring cells prior to the onset of apical constriction (Fig 3D’’; onset of apical constriction is t=0 in Fig 3). Consistently, high proneural index values were measured for *sc* in clusters 2 and 4 from the onset of *sc* expression to NB delamination (Fig 3E’’). In contrast, we observed that the *lsc* gene was detectably expressed only after the onset of NB delamination in clusters 4 (Fig 3D’), and a sharp increase in proneural index values was noted for *lsc* at the onset of apical constriction (Fig 3E’’). This indicated that_the *sc* and *lsc* genes are sequentially activated in clusters 4. It also raised the possibility that cross-regulatory interactions between proneural genes, with Sc positively regulating *lsc* gene expression, contribute to proneural expression dynamics in a PNC-specific manner.

### Deletion of the *AS-C* delayed the apical constriction of presumptive NBs

Since proneural factors were found to accumulate in presumptive NBs prior to their delamination, we wondered whether the proneural genes act to regulate apical constriction. Previous studies using embryos lacking all three proneural genes reported that several VN cells delaminate but fail to express mature NB markers^35^. However, because these experiments relied on fixed samples, it remained unclear whether all presumptive S1 NBs delaminated normally. To address this, we performed live imaging of *Df(1)AS-C* embryos expressing PHmCh to monitor apical constriction and delamination of S1 NBs. Individual embryos were genotyped after imaging by gPCR, using PCR primers to distinguish *Df(1)AS-C* male embryos from wild-type and heterozygous siblings (pooled as controls in Fig 4). Using this approach, we observed no difference in the spatial pattern of NB delamination between *Df(1)AS-C* and control embryos in the medial region of the VN (Fig 4A). To quantify delamination dynamics, we tracked individual NBs from the onset of apical constriction and measured the timing of delamination initiation and the rate of apical surface reduction. We found that *Df(1)AS-C* NBs showed a delay in the onset of apical constriction. While control NBs began delamination 10.8 +/-5.2 min after the start of GBE, *Df(1)AS-C* NBs initiated delamination ∼6 minutes later (17.1 +/-4.0 min after GBE onset; Fig 4B). However, once delamination began, *Df(1)AS-C* NBs proceeded with the same speed of apical constriction as control embryos (Fig 4C). This indicated that proneural genes are required to regulate the timely onset of NB delamination but are dispensable for the process of NB delamination itself.

**Figure 4.**
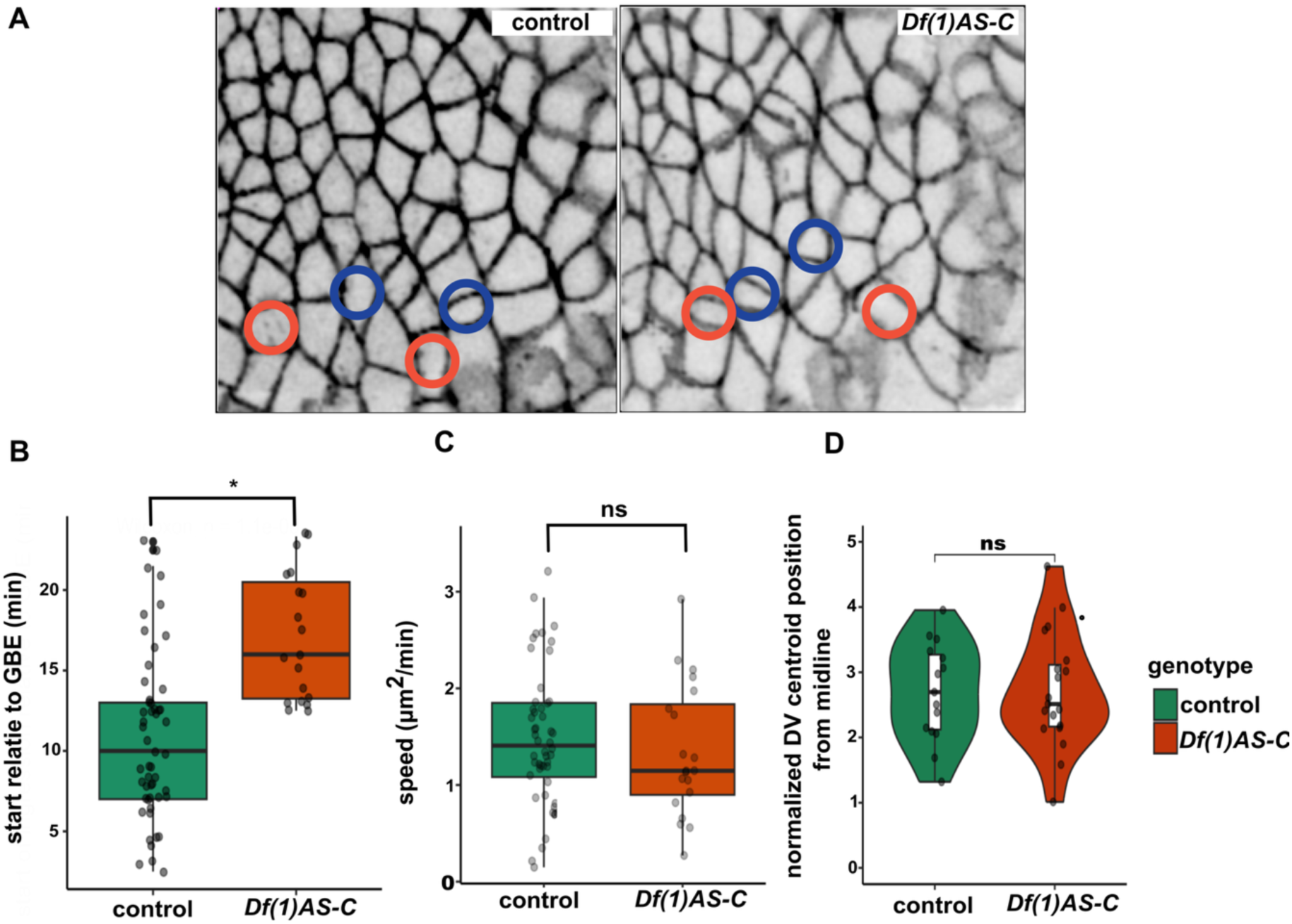
Loss of all proneural genes delayed the onset of NB delamination (A) Representative images of control (left) and *Df(1)AS-C* (right) stage 8 embryos showing delaminating NBs (color-coded as in Fig 1B; PHmCh, black). (B) NB delamination onset relative to GBE in control and *Df(1)AS-C* embryos. (C) Delamination speed in control and *Df(1)AS-C* embryos. Speed was calculated as the slope of area reduction curves. (D) Distributions of the DV positions occupied by presumptive NBs prior to GBE (as in Fig 1E) in control and *Df(1)AS-C* embryos. Violin plots show density; box plots show median and quartiles. In B-D, individual points represent single NBs. *p>0.05, ns = not significant, Wilcox t-test. n = 50 from 4 embryos.

Finally, we used our *Df(1)AS-C* embryo movies to ask whether the DV position bias seen for presumptive NBs prior to GBE (Fig 1D,E) was altered upon loss of proneural activity. Back-tracking medial VN NBs in *Df(1)AS-C* embryo revealed no statistically significant difference between control and proneural mutant embryos (Fig 4D). This indicated that the spatial DV patterning of the medial NBs did not require proneural gene function. This in turn suggested that positional cues act, at least in part, in a proneural-independent manner to establish this spatial bias in NB fate specification.

### NBs initiate delamination before Notch activation in neighboring cells

Notch-mediated lateral inhibition has long been considered a central mechanism in NB specification. Upon loss of Notch signaling, proneural gene expression is not suppressed in neighboring cells and multiple PNC cells delaminate^17,21,25,26,36,37^. However, our observations of an early and strong bias in proneural gene expression in presumptive NBs and of an increase in proneural gene expression in neighboring cells during GBE led us to wonder when Notch signaling is activated to select NBs and suppress proneural gene expression. To measure Notch signaling, we monitored the expression of GFP-m8 using a GFP-tagged knock-in allele of the *E(spl)m8* gene (see Methods), the earliest and strongest direct target of Notch in the VN ^13,25,38^. While GFP-m8 could be detected in fixed stage 8 embryos using anti-GFP, the fluorescence signal was too low to reliably track Notch activity by live imaging, presumably due to the fast turnover of the protein relative to GFP maturation kinetics. We therefore studied Notch signaling dynamics in fixed sample analysis, but this created a challenge: how do we identify NBs without the benefit of time-series data to identify the delaminating cells? To address this problem, we developed a morphometric approach that used an “area index” defined as the cell apical area normalized by the mean area of its immediate neighbors. Using PHmCh embryo movies (Fig 1), we measured the area index of medial VN cells over time and found that area index values decreased over time in NBs as they apically constrict and delaminate while area index values slightly increased in other cells neuroectodermal cells throughout stage 8 (Fig S2A,B). This suggested that this area index could be used to predict NBs. Using these live imaging data, we trained a logistic regression model that predicted NB fate based on both area index and apical area values. This model achieved 75% accuracy in distinguishing NBs from epithelial cells, with a threshold of area index <0.6 (Fig S2C). We applied this NB prediction model to fixed embryos co-stained for E-cadherin (to mark apical cell boundaries) and GFP-m8 (to report Notch activity) (Fig 5A-B’). We observed on average 6 +/- 5 predicted NBs per embryos in early stage 8 embryos (Fig 5A). In mid-stage 8 embryos, we observed 17 +/- 3 predicted NBs per embryo (Fig 5A’,B’). To examine the temporal relationship between NB delamination and Notch activation, we identified NB by area index (model-predicted NBs) and then classified them based on GFP-m8 expression in adjacent PNC cells (PNCs were scored positive when at least one neighboring cell appeared to express GFP-m8; see Fig S2C). This analysis revealed that only ∼5% of the model-predicted NBs had positive neighbors in early stage 8 embryos, whereas this proportion increased to ∼50% in mid-stage 8 embryos (Fig 5C). Additionally, model-predicted NBs within negative PNCs had a mean area index of 0.46 at early stage 8, while those within positive PNCs exhibited a significantly lower area index of 0.38, suggesting that the positive PNCs were more mature than the negative ones (Fig 5D). By mid-stage 8, model-predicted NBs within negative PNCs showed a mean area index like the one measured at early stage 8 (Fig 5D), consistent with the emergence of newly apically constricting NBs. In contrast, the proportion of positive PNCs increased significantly (Fig 5C) and the mean area index of these predicted NBs decreased, consistent with progressive delamination (Fig 5D). To better visualize this progression, we arranged model-predicted NBs in a pseudo-time series ordered by area index values (Fig 5E) and quantified the GFP-m8 signal measured in surrounding cells (Fig 5F, Fig S2E). No significant GFP-m8 expression was detected above normalized signal noise (mean shown as a dotted line in Fig 5F, with the 95% confidence interval in grey) before the NB area index decreased down to a value of 0.4 (Fig 5F). Since *E(spl)m8* is the earliest and strongest effector Notch targets during early neurogenesis^13,25,38^, and since detection of GFP by antibodies is a reliable and sensitive method of immunodetection, our data indicated that NBs delamination precedes Notch receptor activation, at least in some PNCs. Of note, our results are consistent with earlier live imaging of *ms2: E(spl)m8* embryos indicating that the *E(spl)m8* gene was not detectably expressed in the VN up to 30 min after the onset of GBE^39,40^.

**Figure 5.**
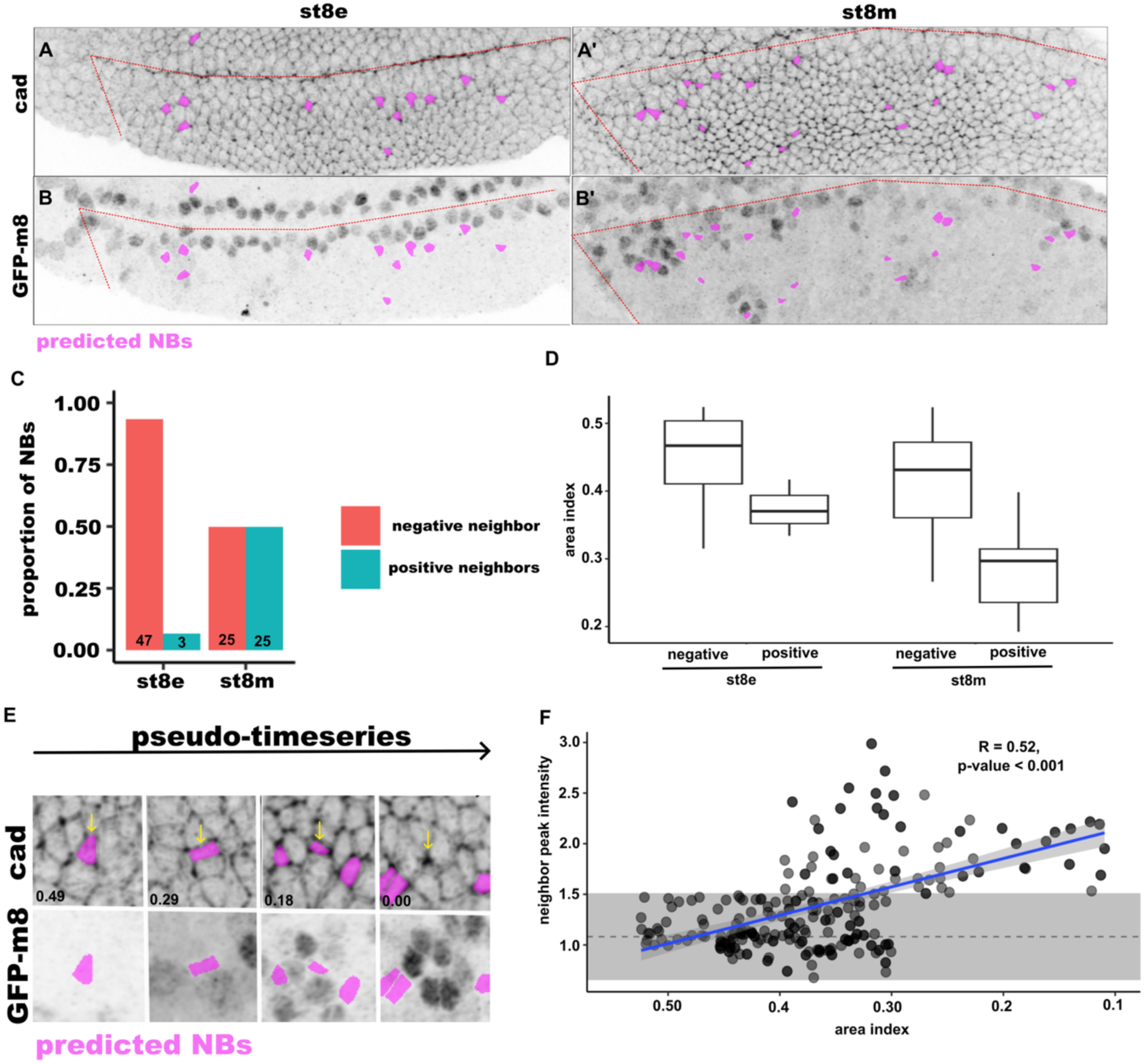
The onset of NB delamination appeared to precede Notch activation. (A-B’) Model-predicted NBs (magenta) in early and mid-stage 8 embryos (noted here as st8e and st8m respectively) immunostained for E-cadherin (A,B) and GFP-m8 (A’B’). Red dotted lines indicate the cephalic furrow and the midline. (C) Relative proportions of model-predicted NBs associated with GFP-m8 negative and GFP-m8 positive PNCs (see Fig S2 for method). In early stage 8 embryos, most model-predicted NBs belonged to PNCs with no detectable GFP-m8 expression. (D) Box plots showing the area index values of the NBs studied in panel C (median and quartiles are shown). (E) Pseudo-time series of model-predicted NBs (magenta) arranged by increasing area index values (top row), from early apical constriction to late delamination (top row: Ecad; bottom row: GFP-m8). (F) Plot of the GFP-m8 signal measured in cells surrounding model-predicted NBs (see Fig S2 for method) as a function of the area index (n=13 embryos; 279 model-predicted NBs). Dotted line represents the mean noise signal (GFP-m8 expression in cells surrounding cells that were not predicted as NBs; the grey area indicates a 95% confidence interval of non-predicted NBs. For all quantifications 10 st8e embryos and 7 st8m embryos were analyzed.

Together, our data clearly indicated that Notch receptor was first activated in PNC cells after NBs have started to apically constrict. This finding has important implications about the role of Notch in NB selection and delamination, as discussed below, leading us to revise the current model of NB fate selection.

## Discussion

Quantitative live imaging of *lsc* and *sc* transcription revealed early spatial biases in proneural activity along the DV axis. It also showed that initial differences in gene expression are maintained from the earliest detectable expression through the onset of NB delamination. Thus, these initial biases were found to predict NB fate acquisition. Additionally, Notch activity within PNCs, as measured by *E(spl)m8* expression, was not detectable until after NB delamination had initiated. Together, these results implied that early NB specification and delamination initiation occur prior to, hence independently of, Notch signaling. Our results therefore challenge current models whereby lateral inhibition singles out NBs from within groups of equipotent cells through stochastic symmetry breaking or via the amplification of an initial bias by Notch^41^. In contrast with these models, we propose that positional cues bias early proneural gene expression and direct early NB fate acquisition whereas Notch acts later to stabilize and reinforce these initial fate differences by suppressing the proneural potential in other PNC cells. Notch would also act to prevent the delamination of these PNC cells. In this model, NBs are pre-defined by positional cues in the embryo and Notch does not single out NBs from PNCs (Fig 6A).

**Figure 6.**
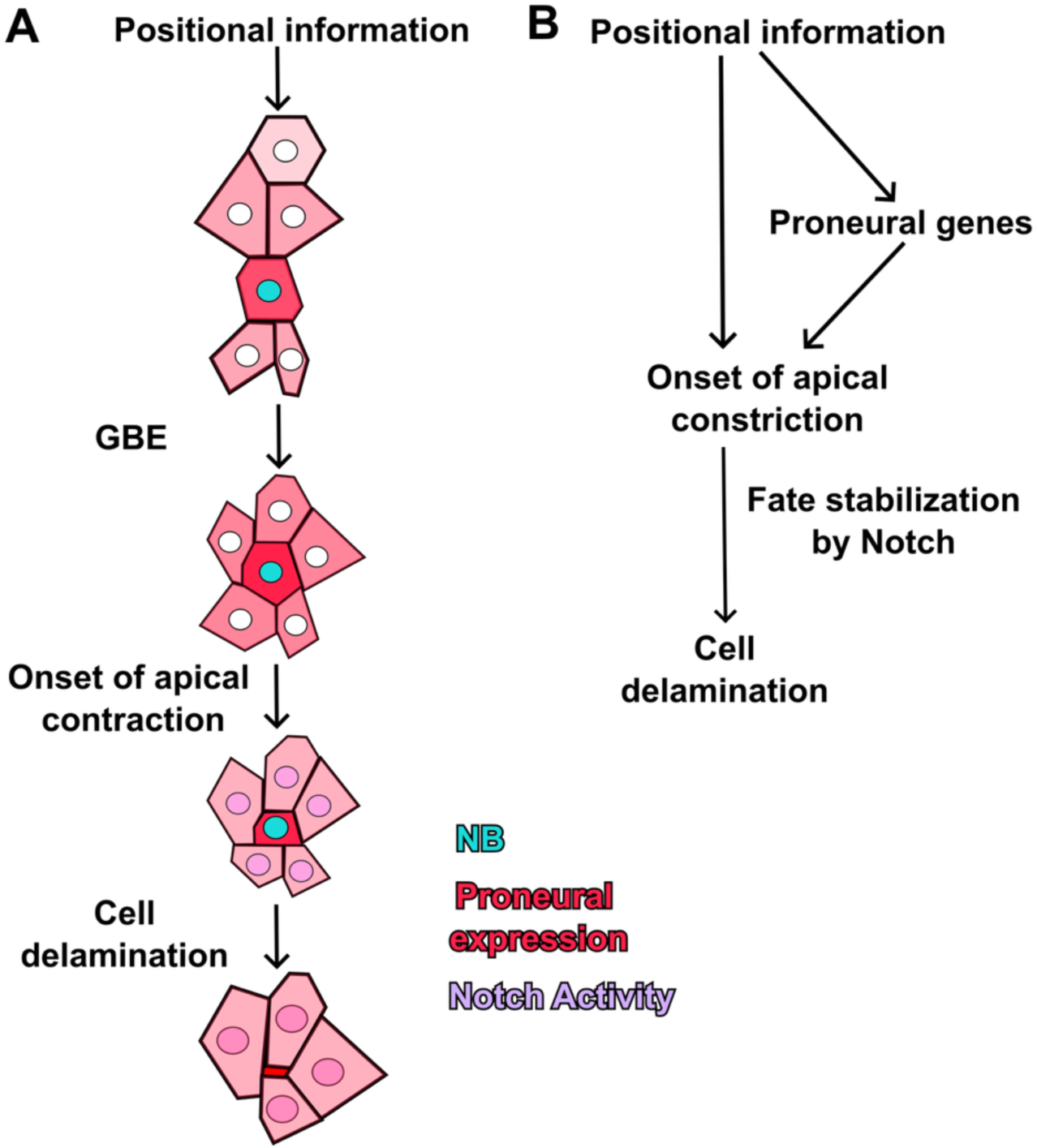
Model (A) Positional cues provide a DV bias for early proneural expression (intensity color-coded in red). During GBE, NBs (nucleus in cyan) apically constrict. The onset of delamination precedes Notch receptor activation (magenta) in neighboring PNC cells. (B) Positional cues are proposed to regulate the onset of apical constriction in individual cells both directly and via the regulation of the proneural genes. During delamination, Notch is proposed to stabilize fate choices.

This model is in part based on our ability to detect early Notch activity in PNCs. Here, we used endogenous GFP-E(spl)m8 for the following reasons. First, the *E(spl)m8* gene appears to be amongst the first *E(spl)-C* gene to be expressed at high levels in the VN^38^. Second, since the E(spl) proteins mediate the function of Notch in restricting NB fate acquisition to a single cell per PNC^42,43^, E(spl) protein accumulation can serve to report when Notch signaling becomes active. Third, very low levels of GFP-m8 proteins can be detected using high-affinity anti-GFP antibodies in fixed embryos. Combining this sensitive approach with a model that predicts NBs, we showed that 94% of predicted NBs in early stage 8 embryos had no detectable GFP-m8 expression in neighboring cells. Given our model’s 75% accuracy threshold, and conservatively assuming all prediction errors are false positives, 70% of all PNCs appear to lack *E(spl)m8* expression prior to the onset of NB delamination. Thus, Notch does not appear to act for the initial selection of NBs from PNCs. A similar conclusion is reported in a companion paper from the Bray lab who used live imaging of *E(spl)m8* transcription to show that it is never detected in presumptive NBs but only in surrounding cells. They further propose that there is already an apical size bias when *E(spl)m8* is first transcribed, in line with it occurring after NB selection has been initiated. Thus, Notch signaling may primarily function during early neurogenesis to amplify pre-existing differences and stabilize cell fates by continuously reinforcing initial fate differences through feedback, hence providing robustness against perturbations. We propose that positional information, acting in part via the proneural genes, pre-selects individual cells to become NBs. More generally, the extent to which cell fate choice during lateral inhibition involves the amplification of stochastic fluctuations of Notch via an intercellular feedback loop, as proposed earlier^10^, or relies on early cell-to-cell differences in proneural gene expression with Notch acting later to stabilize cell fate remains to be determined^9^.

How spatial biases in early proneural gene expression are established is not known. Previous studies have indicated that the precision of positional information along the AP axis is sufficient for each cell to have a unique cell identity^44^. Therefore, AP patterning factors could potentially direct the row-specific expression of the *lsc*, *ac* and *sc* genes (Fig 1B)^17^. In contrast, how positional cues could bias proneural gene expression along the DV axis is less clear. However, previous studies have shown that EGF receptor (EGFR) signaling provides positional information to pattern the ventral ectoderm^45,46^ and neuroectoderm^47^, acting for instance to pattern the VN into columns via the regulation of the *vnd* and *ind* genes^48^. Whether EGFR signaling can provide DV patterning cues at a finer spatial resolution within individual PNCs remains to be studied.

Interestingly, we observed that the spatial pattern of delaminating cells persists in the complete absence of proneural activity, indicating that positional cues may be sufficient to define a regular pattern of delaminating cells. This in turn suggests that the onset of delamination is regulated by both proneural factors, shown here to promote the onset of apical constriction, and by upstream regulators that may provide positional information for both early proneural factor activity and cell delamination (Fig 6B). Of note, the extreme speed at which early embryogenesis proceeds in *Drosophila* may favor the selection of such feedforward regulatory mechanisms whereby transcription factors that act early to pattern the embryo and that are still present during GBE can also serve to pattern cell fate during neurogenesis. Likewise, the key role of positional cues in NB selection might reflect temporal constraints on NB specification in the fast-developing *Drosophila* embryos. While Notch was initially proposed to select individual cells from within equipotential groups, it has become clear that Notch operates along a spectrum, with varying degrees of positional information guiding fate outcomes^13,41,48,49^. NB selection in the early embryo lies at one end of this spectrum, with positional information having an early and predominant role in selecting NBs and Notch acting later to stabilize an initial difference in cell fate. Thus, the integration of positional information with Notch signaling is context dependent, with developmental speed and tissue architecture possibly influencing the relative contributions of self-organization versus positional information to generate stereotyped cell fate patterns.

## Acknowledgements

We thank Y. Bellaiche, M. Laghia, Bloomington Drosophila Stock Center (BDSC) for flies, Flybase for database services, Addgene for plasmids and the Developmental Studies Hybridoma Bank for antibodies. We thank L. Couturier for help in generating the *E(spl)m8^GFP^* and *lsc^GFP^* lines. Thank B. Zoller and T. Gregor for help with analysis of MS2 spots. We are grateful to R Levayer and all lab members for discussion and critical reading. This work was funded by grants from the Agence Nationale pour la Recherche (ANR-10-LABX-0073) and the Fondation pour la Recherche Médicale (FRM-DEQ20180339219) to FS.

## Materials and Methods

### Flies

The following lines were used: ubiP-PLCγPH-ChFP (noted here PHmCh)^50^; *nosP-MCP-eGFP, ubiP-His2Av-mRFP* (which produce MCP-GFP and His2A-RFP)^51,52^; *Df(1)BSC530* (BL-25058; this 259 kb deletion, noted *Df(1)AS-C* herein, deletes the entire *AS-C*). All strains were maintained on standard cornmeal medium at 25°C.

### Genome engineering

The *lsc* gene was GFP-tagged to produce the *lsc^GFP^*line using Cluster Regularly Interspaced Short Palindromic Repeats (CRISPR)-mediated Homologous Recombination (HR) using the same gRNAs as for *sc^GFP^*^49^. To do so, Cas9-expressing embryos were injected with a mix of plasmids encoding gRNAs flanking the region to be recombined together with a donor template carrying a 3xP3-RFP selection marker flanked by loxP sites. The donor template for HR was first produced by BAC recombineering in *E. coli* and then transferred into a multi-copy vector as previously described^53^. The BAC CH322-63B05 encoding the *lsc* gene was used to introduce sfGFP flanked with a GVG linker at the terminus of the ORF. The 3xP3-RFP selection marker flanked by loxP sites was produced by gene synthesis. The *E(spl)m8* gene was GFP-tagged using a similar approach. A BAC transgene encoding the GFP-tagged E(spl)m8 protein, described in Trylinski *et al.* (2017)^54^, was used to generate a multi-copy donor plasmid using BAC recombineering. This donor plasmid was used to perform CRISPR-mediated HR to tag the endogenous gene at the *E(spl)-C*. The 3xP3-RFP selection marker flanked by loxP sites was used to select the knock-in line. The resulting *E(spl)m8^GFP^* line is deposited at VDRC (#311302). For *lsc^GFP^* and *E(spl)m8^GFP^*, a mix of donor template (300 ng/μl) and gRNA plasmids (150 ng/μl) was injected into 900-1,200 embryos from the PBac{vas-Cas9}VK00027 stock (BL-51324).

The *lsc^ms^*^2^ *and sc^ms2^* knock-in lines were produced by CRISPR-HR using the same gRNAs as those used for *lsc^GFP^* and *sc^GFP^*, respectively. The 24xMS2 cassette stem loop repeats block was obtained by digestion of the pCR4-25MS2SL-stable plasmid (Addgene 31865) and recombined with the same donor template plasmids as *lsc^GFP^* and *sc^GFP^* replacing their sfGFP. The design of the final donor plasmid was such that the MS2 block is located 7bp upstream of the ATG start codon and is flanked by the 5’ and 3’ parts of the 55bp intron of the *Rpl5* gene (this intron was chosen because it was small and efficiently spliced from an abundantly expressed gene). For each of these CRISPR knock-in lines, HR at the locus was verified by genomic PCR. For the *lsc^ms2^ and sc^ms2^*knock-in lines, injection was performed by BestGene (Chinmo, US). Cloning details and gRNA sequences are available upon request.

### Live imaging

For live imaging experiments, males carrying the *ms2:sc* (or *ms2:lsc*) alleles were crossed to MCP-GFP, His2A-RFP; PHmCh females to generate embryos with the following genotype: *ms2:sc* (or *ms2:lsc*)/+; MCP-GFP, His2A-RFP /+; PH-mCh/+. Stage 6 (pre-gastrulation) embryos were dechorionated in 50% bleach for 2 min, rinsed extensively in water and mounted ventral side down in Halocarbon oil 27 (Sigma-Aldrich). Embryos were transferred to a gas permeable membrane (YSI Life Sciences) prior to imaging at 25°C.

Image acquisition was performed on a Nikon eclipse Ti2 inverted confocal microscope using a 40X Plan Apo silicone oil immersion lens (NA 1.25, Nikon) using a 488nm laser (6.5% power) and a 561nm laser (3% power). Images were acquired using NIS-Elements software (Nikon). For apical surface tracking z stacks of 7 um (8 slices Δz=1um) were acquired every 15 seconds for 60-90 min. For experiments tracking both apical surface and nuclei, z-stacks of 30um (31 slices, Δz=1um) were acquired every 30 seconds. All images shown represent maximum intensity projections unless otherwise noted.

### Image analysis and quantification

#### Segmentation/Tracking of MS2 spots

All image analysis was performed in Python using custom scripts (available at https://gitlab.pasteur.fr/4dunit). Images were pre-processed by selecting relevant z-planes for each channel: single apical surface plane was manually identified for each timepoint (to correct for z drift) to track apical movements, while all planes containing nuclear signal were retained for 3D nuclear segmentation.

Apical cell surfaces were segment in 2D using Cellpose (v3.1, cyto3 model)^55^. Cells were tracked across time using Epicure plugin in Napari^56^. Nuclei were segmented in 3D using Cellpose (v3.1, deblur_cyto3 model) and tracked using TrackMate^57,58^.

#### MS2 spot detection and intensity measurement

MS2 transcriptional spots were detected in the GFP channel using intensity-based thresholding (threshold = mean + 3x SD of background fluorescence) for each nuclear volume. For each tracked nucleus at each timeframe, we identified whether a thresholded spot was within the 3D volume of the nuclear mask. When present, we measured total GFP fluorescence intensity with a cylindrical volume (radius 6px, height 3px) centered on the spot. Background fluorescence was calculated as median intensity in an annular region (inner radius = 6px, outer ring = 8px) around each spot and subtracted. Spot intensities were normalized to background fluorescence to control for expression level variation of MCP-GFP and affects from nuclear depth.

**Cumulative intensity** for each cell was calculated as the sum of background-corrected spot intensities across all timepoints.

**Proneural index** is calculated as

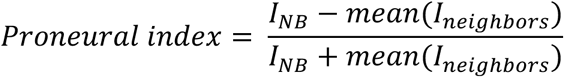

where I_NB_ is the cumulative spot intensity in the presumptive NB and I_neighbors_ is the mean intensity of all cells sharing a contact with the NB at the apical surface as determined by Epicure^56^.

#### NB identification and alignment

NBs were retrospectively identified by their characteristic apical contraction and delamination^26^. For temporal analyses the onset of delamination was defined as the first frame where the apical surface begins to decrease below 40um with no significant rebound. The onset of GBE was identified by the first frame where posterior flow of the ventral ectoderm can be observed.

#### Apical surface segmentation

For tracking NB delamination and fixed samples were apical surface measurements were needed images were processed using max z-intensity projections for the most apical 2 z frames. Segmentations were performed automatically using Epyseg^59^ and corrected with Epicure^56^.

#### Quantitative analysis of the GFP-m8 signal

To quantify GFP-m8 levels in PNC cells surrounding model-predicted NBs, we used our apical surface segmentations to identify the centroid of each cell and mapped it to the corresponding position on the GFP-m8 channel (Fig S2D). We then generated a series of concentric rings around the centroid and measured the fluorescent intensity within each annulus (Fig S2E). We then calculated the mean of 3 annuli intensity around the peak intensity normalized to the background of the entire image.

### Fixed sample preparation

Embryos were dechorionated and fixed following standard methods. The following antibodies were used: rat anti-Ecad (DCAD2 from DHSB, 1:100), goat anti-GFP (Abcam, ab6673, 1:1000) and mouse anti-En (4D9 from DHSB, 1:1000). Secondary antibodies were from Jackson ImmunoResearch: anti-goat Alex 488(1:400, 705-545-003), anti-rat Cy3 (1:400, 712-165-150) and anti-mouse Cy3 (1:400, 715-165-150). Images were acquired with Nikon Eclipse Ti2 inverted microscope with a 40X Plan Apo silicone oil immersion lens (NA 1.25, Nikon).

### Genotyping *Df(1)AS-C* embryos

The BSC530 deficiency was generated using FLP-mediated recombination between two mapped FRT sites^60^. This recombination event had left elements of the transposable element at the site of the deletion, and primers had been previously designed to screen for the presence of these elements^60^. In addition, we generated primers that span the 3’ deficiency breakpoint. After imaging, individual embryos were crushed in DNA extraction buffer (100 mM Tris-Cl, pH 8.2, 1 mM EDTA, and 25 mM NaCl) and prepared for single embryo PCR following standard methods^61^. Embryos that were positive for the transposable element, but negative for the breakpoint primers were determined to be mutant *Df(1)AS-C* /Y embryos.

### NB Identification Model

Individual frames from live imaging data of PHmCh embryos obtained 20-40 min post-GBE onset were split 80/20 into training and testing sets. We trained a logistic regression model using area and area index as predictors with stats::glm (v4.5.2) function in R. Area index was calculated as the area of the cell of interest normalized by the mean area its neighbors, defined as cells with direct cell-cell contacts as determined by Epicure. Model performance was evaluated using F1 score, which represents the harmonic mean of precision (susceptibility to false positives) and recall (susceptibility to false negatives). Classification thresholds were systematically tested to identify the optimal boundaries.

**Movie 1 – Cell intercalation of ventral VN during GBE** – related to Fig 1C. PHmch embryo during GBE (right). Segmented and tracked VN (red) and mesoderm (blue) (left). Time in min:sec.

**Movie 2 – *lsc:MS2* expression dynamics** – related to Fig 2B. Live time series during NB specification. Phmch (grey) and MCP-GFP (*lsc* transcription) (green). Time in min:sec.

**Movie 3 – *sc:MS2* expression dynamics** – related to Fig 2B. Live time series during NB specification. Phmch (grey) and MCP-GFP (*sc* transcription) (green). Time in min:sec.

**Movie 4 – *lsc:MS2* DV expression.** Expression of *lsc* along DV column pseudocloured by cumulative proneural expression measured by MCP-GFP. Darker colors represent low expression and brighter colors high expression. Time in min:sec.

**Movie 5 – *sc:MS2* DV expression.** Expression of *sc* along DV column pseudocloured by cumulative proneural expression measured by MCP-GFP. Darker colors represent low expression and brighter colors high expression. Time in min:sec.

---

**Figure S1.**
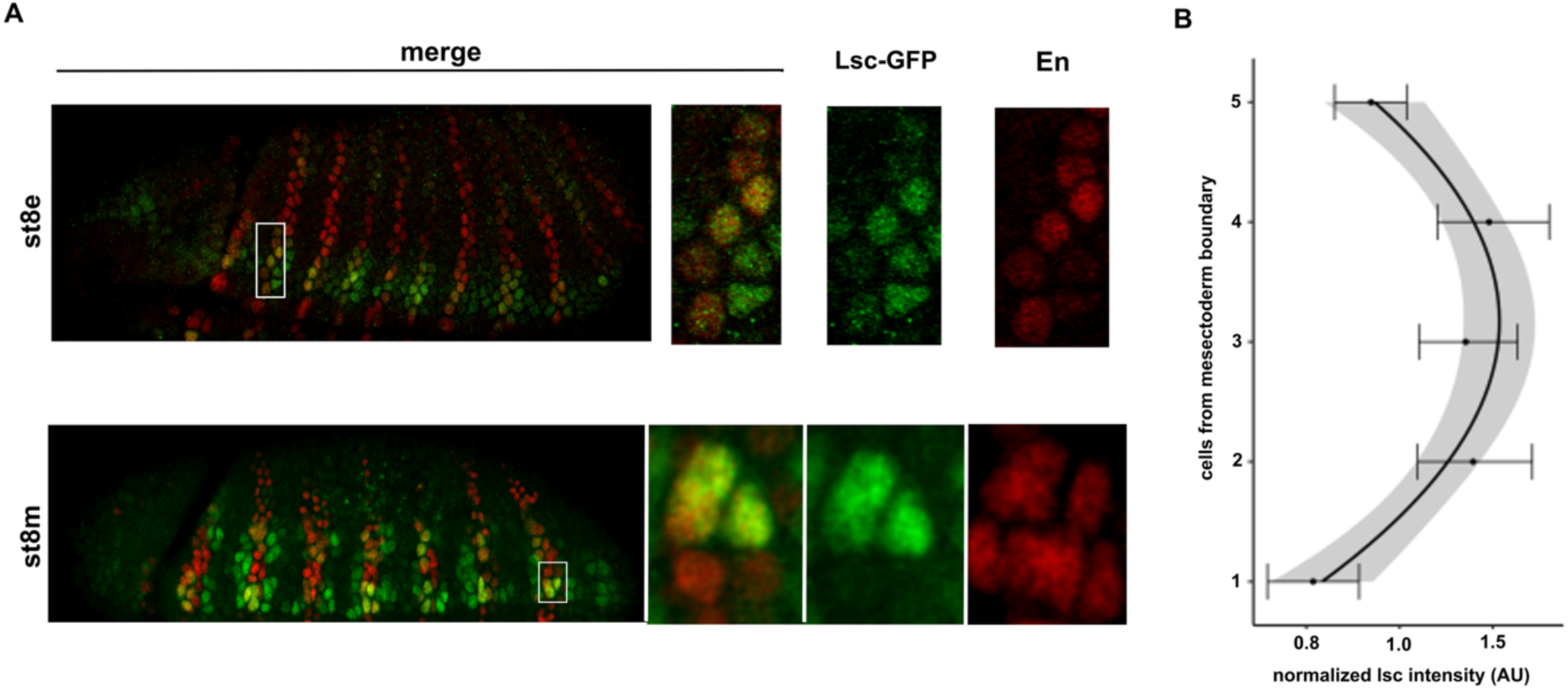
: An early DV bias in Lsc expression (A) Lateral views of early and mid-stage 8 embryos showing the distribution of the Lsc protein (GFP-Lsc, green; Engrailed (En), red) in low- and high magnification views. (B) The analysis of normalized Lsc intensity along the DV axis showed higher levels of Lsc protein in cells located 2-4 cells away from the midline. Grey region represents standard deviation (n= 25 PNCs from 7 embryos).

**Figure S2:**
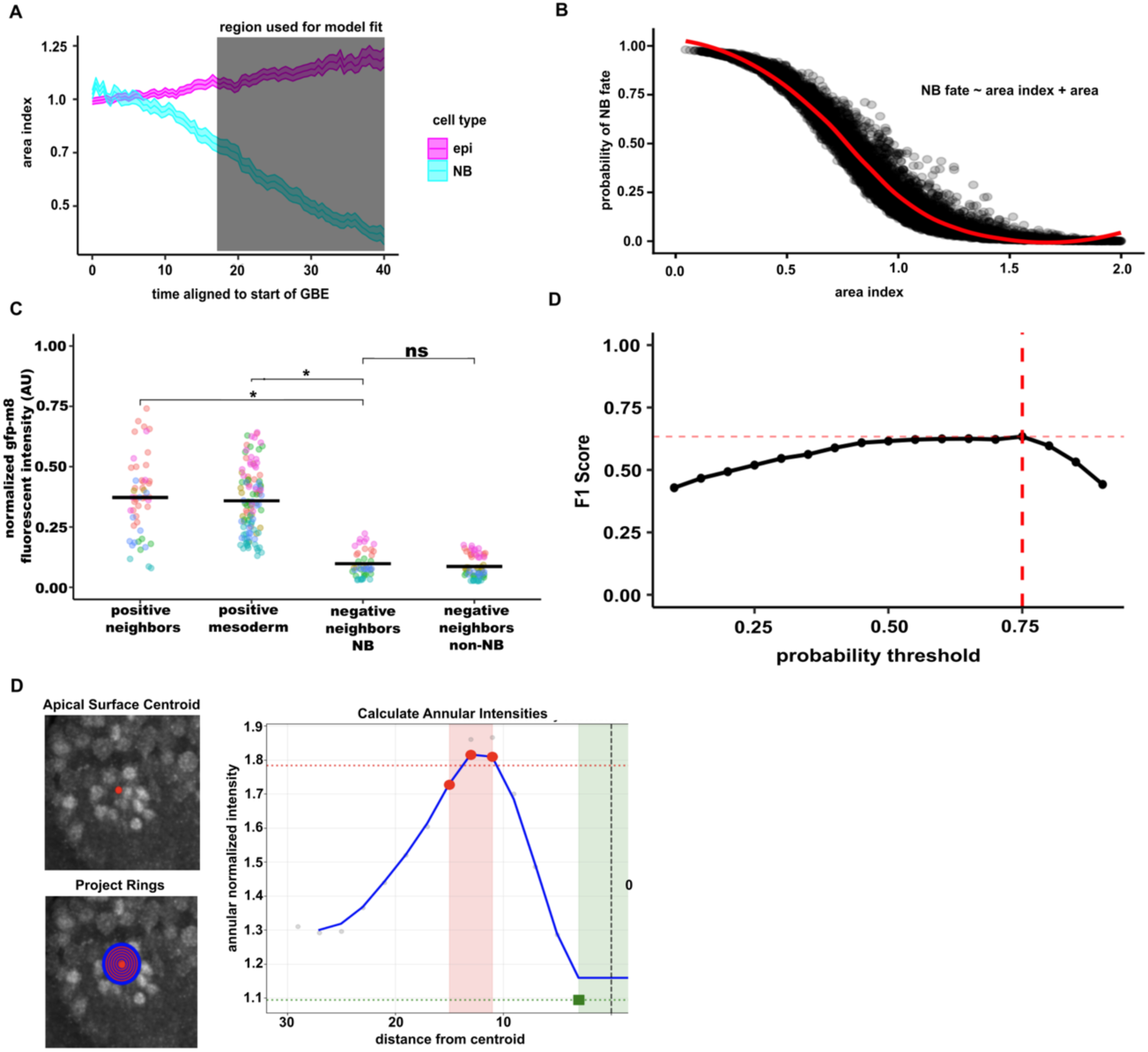
Quantitative modelling methods for fixed sample analysis. (A) Temporal dynamics of the area index for epithelial cells (epi, magenta) and NBs (cyan) aligned to the onset of GBE measured from PHmCh movies. Grey shaded region indicates time window used for model fitting. n=5 embryos. (B) Logistic regression model predicting NB fate based on area index and area size. Each point represents one cell from one frame. (C) Normalized GFP-m8 expression in manually categorized cell populations from fixed stage 8 embryos: GFP-m8⁺ cells from positive PNCs, GFP-m8⁺ mesectodermal cells, cells from negative PNCs (adjacent to predicted NBs or non-NB cells). Each dot represents the value from a single nuclear ROI; colors denote individual embryos. Wilcoxon test, * p < 0.05, ns = not significant, n=6 embryos, ∼50 ROIs (i.e. nuclei) per category. (D) Model optimization: F1 score versus probability threshold for cell fate classification. Red dashed line indicates the 0.75 probability threshold that was used to identify NBs in fixed samples. (E) Schematic of annular quantification method. Concentric rings projected from apical centroid for spatial intensity analysis. Normalized intensity profiles plotted versus distance from centroid. Mean peak intensity quantifies neighbor intensity

## Notes

### Competing Interest Statement

The authors have declared no competing interest.

