## Supplementary figures and images for "Positional cues, not Notch, direct Neuroblast selection during early neurogenesis in the *Drosophila* embryo"

### Supplemental Figure 1

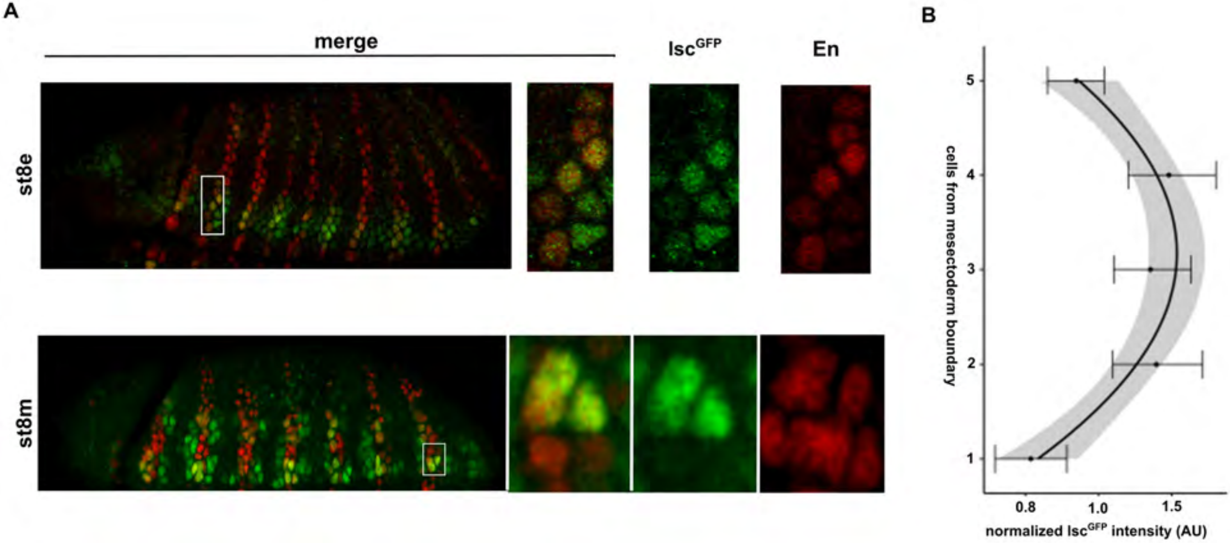

### Supplemental Figure 2

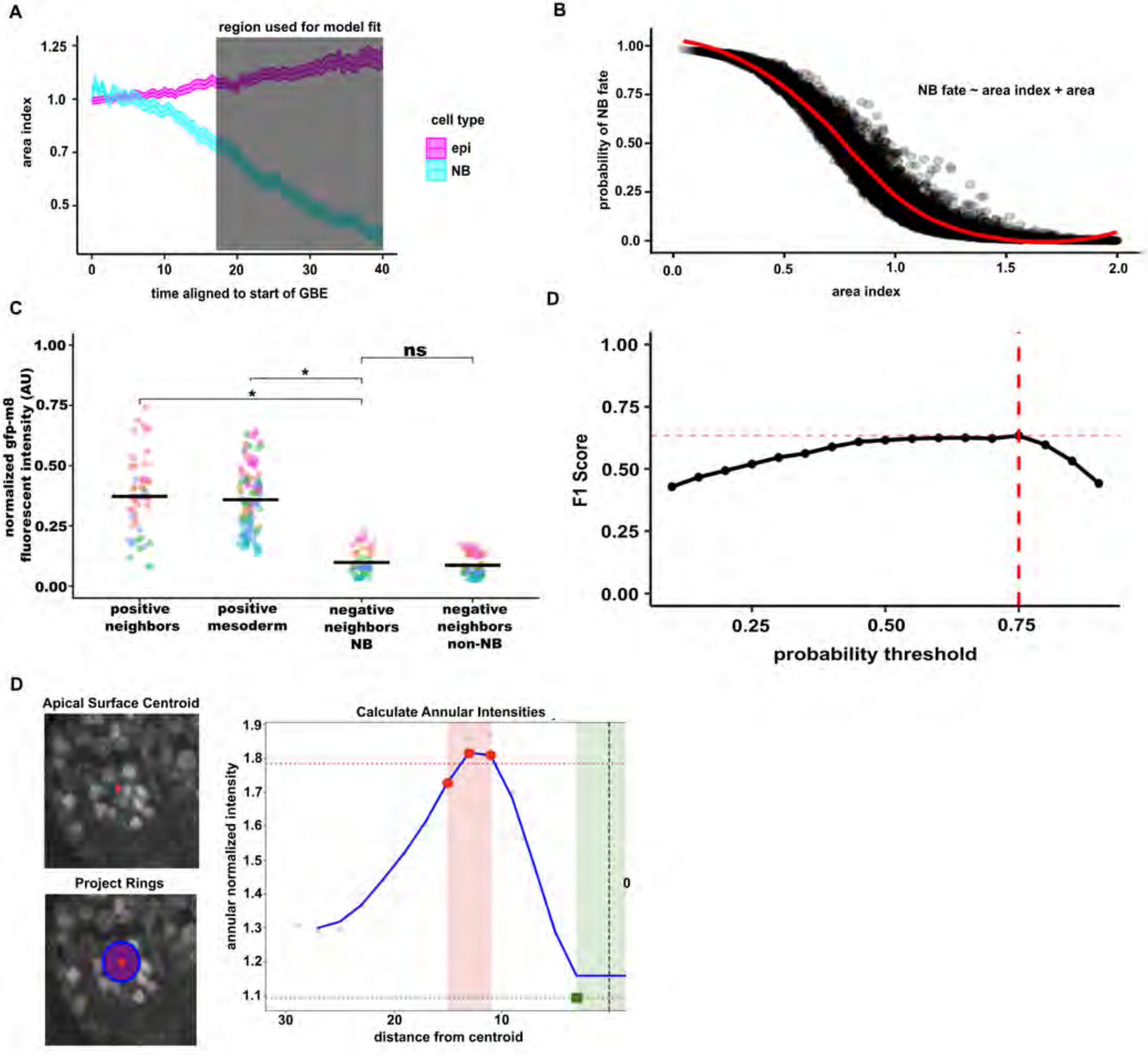
